# Women and Men Were Proportionally Represented Among Speakers at Major National Neurology Conferences in 2017

**DOI:** 10.1101/866160

**Authors:** Mollie McDermott, James F. Burke, Haley McCalpin, Anita V. Shelgikar, Douglas J. Gelb, Abbey Dunn, Nicholas J. Beimer, Zachary N. London

## Abstract

**Objective:** To determine whether speaking roles at five major neurology conferences in 2017 show disproportionate representation by men.

**Methods:** This study consisted of two cross-sectional analyses. In the first part, we compared speaker characteristics across meetings and by gender using descriptive statistics. In the second part, we linked presenters to the American Medical Association (AMA) Masterfile. For the primary analysis in the second part using linked AMA speaker data, we built models to estimate the influence of gender on speaker roles.

**Results:** 1493 speakers were identified and included in our cross-sectional analysis. Women made up 28% of presenters from the US and 18% of presenters from other countries. After adjusting for years from medical school graduation and subspecialty, no effect of gender on speaker activity was observed (odds ratio [OR] for women 0.91; 95% confidence interval [CI], 0.77-1.07).

**Conclusions:** Factors aside from national conference speaking activity should be investigated to better understand sex differences in rank at top-ranked academic neurology programs.

## Introduction

Gender equity in academic neurology is a topic of growing interest. A 2018 study of 29 top-ranked academic neurology programs found that men outnumber women at all faculty ranks, and this discrepancy increases with advancing rank.^1^ The reasons that fewer women hold senior academic ranks are unknown. In the past, more men than women entered neurology (and medicine in general), but the overrepresentation of men at senior ranks in academic neurology persists even after correcting for the relative proportions of men and women in the overall population of neurologists. Men have more publications than women at each rank, but other measures of academic productivity including clinical activity, educational leadership, and book authorship do not differ between men and women.^1^ One possible driver for the gender disparity in academic neurology promotion is participation as a speaker at national conferences. In many academic neurology departments, lecturing at national conferences is an important factor in academic promotion.

Some literature exists regarding the question of gender disparity in other specialties’ conferneces, but the conclusions vary. For example, a 2018 retrospective analysis of scientific programs from three international and two national critical care annual conferences from 2010 to 2016 found that female physicians represented 5-26% of speakers, while 30-35% of clinical care trainees at that time were women.^2^

In contrast, an analysis of programs from eight international emergency medicine conferences from 2014 to 2015 found that only 29.9% of presentations were given by women.^3^ However, the proportion of qualified female emergency medicine specialists in the United States, Canada, United Kingdom, and Australia ranged from 23.5 to 30.0%, suggesting that no relative discrepancy exists. Supporting these findings, a recent large analysis of 181 medical and surgical conferences in the US and Canada found that over the last decade (2007-2017) the mean proportion of female speakers (34.1%) was similar to the mean proportion of active female physicians across all specialties (32.4%) during the same time period.^4^

Women may be less likely to present specifically in invited plenary sessions at national conferences. An analysis of plenary speakers at the 2016 European Association of Palliative Care conference found that, although 73.8% of speakers were women, only 6 of 23 (26.1%) invited plenary speakers were women.^5^

The gender distribution of speakers at national neurology conferences is unknown. We used publicly available data to investigate this issue. Given the gender disparities observed in academic neurology rank^1^, we hypothesized that speaking roles at five major neurology conferences would show disproportionate representation by men.

## Methods

We performed two studies: 1) a retrospective cross-sectional analysis of speakership at five major academic neurology conferences in 2017 and 2) a cross-sectional study of all neurologists linked to data regarding speaking roles. Our sought to determine the representation of women as speakers and to explore factors associated with speaking activity.

### Identification of Conferences

The authors queried academic neurology colleagues about the most popular North American conferences in their respective subspecialties. After collecting a comprehensive list, the number of attendees at each conference was recorded (Table 1). Conference attendee numbers were collected through online research or direct calls to the conference governing organizations and reflect either averages or exact numbers from the 2017 annual conferences. The conferences were then ranked based on attendance, and the five largest U.S. conferences were selected for further analysis. All data were publicly available.

**Table 1.** Largest North American neurology conferences in order of attendance. Attendance numbers are based on averages or most recent annual conference attendance.

|  | **Conference Name** | **Estimated # of Attendees** |
| --- | --- | --- |
| **1** | American Academy of Neurology | 14,000 |
| **2** | International Stroke Conference | 5,611 |
| **3** | Sleep | Just over 5,000 |
| **4** | American Epilepsy Society | 4,000 |
| **5** | Society for Neuro-Oncology | 2,500 |
| **6** | Consortium of Multiple Sclerosis Centers | 2,200 |
| **7** | Child Neurology Society | Just over 2,100 |
| **8** | Neurocritical Care Society | Just over 1,000 |
| **9** | American Neurological Association | 900-1,100 |
| **10** | Americas Committee for Treatment and Research in Multiple Sclerosis | 980 |
| **11** | American Headache Society | 900 |
| **12** | American Association of Neuromuscular and Electrodiagnostic Medicine | 877 |
| **13** | American Pain Society | 720 |

### Identification of Speakers

The 2017 programs from the five conferences were obtained. From each program, the following information was recorded for each presentation: last name of presenter, first name of presenter, home institution, and whether the member was in a leadership position (e.g., conference chair, session chair). Speakers giving oral presentations, platform presentations, invited presentations, honorific lectures, and/or plenary presentations were included. Moderated poster sessions were not included. An Internet search for each presenter was performed to obtain rank, medical graduation year, and whether he or she was in academics or private practice. Finally, using Scopus (http://www.scopus.com), an author search was performed cross-referencing the faculty member’s name and institution. The number of published documents and h-index were recorded.

### Linkage to AMA Physician Masterfile

To further characterize speakers and to understand the population of potential presenters, we linked presenters to the American Medical Association (AMA) Masterfile, a database of all individuals who enter medical or post-graduate training in the United States.^6^ Direct identifiers were not available to link datasets, so we used probabilistic matching with the python record linkage toolkit^7^ to match AMA Masterfile records with individuals in the speaker dataset. Records were matched based on first name, last name, middle initial, gender, specialty, and year of medical school graduation. Records were retained if they matched perfectly across all criteria or if there was substantial separation in match quality between the best possible match and next best possible match. Subjective manual review of the first 100 matches suggested that the overall match quality was high. We were able to link 83% of speakers to the AMA Masterfile. The majority of those who were not linked, after manual review of their data, had some foreign training and/or appeared not to be clinically active.

### Statistical Analysis

In the first part of our study, we compared speaker characteristics across meetings and by gender using descriptive statistics in the speaker dataset. Differences in characteristics were assessed using t-tests for continuous variables and chi-square tests for categorical variables in gender-based comparisons and linear regression and ANOVA, respectively, for meeting-based comparisons.

For the primary analysis in the second part of our study using the linked AMA speaker data, we built models to estimate the influence of gender on speaker roles. Unadjusted models predicted speaker positions as the dependent variable in a logistic regression model. Adjusted models added covariates (specialty and years of training) as independent variables. Year of graduation was treated as a linear term Secondary analyses repeated these models using speaker activity at each individual meeting as the dependent variable. Exploratory analyses modeled the dependent variable as a count of the number of presentations given at a given meeting using Poisson regression models. Finally, within presenters at the American Academy of Neurology (AAN) meeting (the largest meeting in our dataset), we explored factors that predicted speaker roles when adjusting for publication count, Scopus h-index, and whether a speaker has an academic appointment. Using a logistic regression model, we also explored whether speakers were in a leadership position at the AAN. All analyses were performed before and after excluding international speakers, given that we were primarily interested in United States academic neurologists. Analyses were performed using the open source python stats model library.^8^

## Results

Across the five conferences, 1493 speakers were identified and included in our cross-sectional analysis. The total number of speakers and the proportion of those speakers who are women are shown in Table 2. Across all meetings in our analysis, women made up 28% of presenters from a practice or institution in the United States and 18% of presenters from other countries. In comparison, 30% of neurologists in the AMA Masterfile were women and 25% of the linked records were women. The International Stroke Conference (ISC) had the lowest proportion of female speakers at 22% and the AAN had the highest proportion at 31.6%.

**Table 2.** Number of speakers, number and percentage female, average medical school graduation year of speakers, and average h-index of speakers by conference.

| **Conference** | **Number of speakers/**<br>**Number of speakers with gender identified** | **Number (%) Female** | **Average Medical School Graduation Year of Speakers** | **Average h-index of Speakers** |
| --- | --- | --- | --- | --- |
| American Academy of Neurology | 512/512 | 162/512 (31.6) | 1992 | 27.9 |
| International Stroke Conference | 456/420 | 91/420 (21.7) | 1994 | 30.1 |
| Sleep | 145/145 | 44/145 (30.3) | 1991 | 26.4 |
| American Epilepsy Society | 274/271 | 85/271 (31.4) | 1993 | 24.3 |
| Society for Neuro-Oncology | 106/104 | 29/104 (27.9) | 1995 | 37.2 |

In unadjusted analysis in the linked dataset, women were more likely to be speakers at a major national meeting (odds ratio [OR] 1.18; 95% confidence interval [CI], 1.01-1.38). However, after adjusting for years from medical school graduation and subspecialty, no effect for gender was observed (OR 0.91; 95% CI, 0.77-1.07). Similarly, after adjustment, no significant effect of gender on speaker activity was observed for any individual meeting. For the AAN-specific analyses, there were no significant effects of gender on likelihood of being a speaker, number of speaking roles, likelihood of being a conference leader, or number of leadership roles.

## Discussion

In this study we found that while women were the minority of speakers at large U.S. neurology conferences in 2017, women’s representation as speakers was proportional to their representation in the field of neurology. In an analysis limited to the largest conference, there were no significant effects of gender on likelihood of being a speaker, number of speaking roles, likelihood of being a conference leader, or number of leadership roles.

These results failed to confirm our hypothesis that there would be an over-representation of men in speaking roles at major neurology conferences. We had reasoned that the factors influencing the selection of speakers at national conferences would strongly correlate with the factors influencing academic appointments and promotions at top-ranked academic neurology programs, where gender disparity has been shown to exist.^1^ We reasoned that this correlation would apply both to factors related to scholarly productivity and to factors related to explicit or unconscious bias.

The most optimistic interpretation of our findings is that the gender distribution at national neurology conferences fails to mirror the gender distribution in academic neurology departments because conferences can pivot more quickly than academic departments in response to the changing demographics of neurology residency graduates. According to this interpretation, the current gender distribution among speakers at major neurology conferences is a harbinger of more proportionate gender representation in top-ranked academic neurology departments in coming years, once the inertia of the promotions process has been overcome.

A less optimistic interpretation is that although neurology conferences readily reflect the changing demographics of practicing neurologists, academic neurology programs intrinsically lack that flexibility. Alternatively, it could be that neurology conference speaker invitations and academic neurology promotions do not correlate because they recognize different aspects of scholarship or measure scholarship differently.

Regardless of the ultimate implications for academic neurology appointments and promotions, we are encouraged by our discovery that – contrary to our hypothesis – women are proportionally represented among speakers at major national neurology conferences. Nonetheless, women speakers are still underrepresented in proportion to the number of women entering the field of neurology. In 2017, 44.7% of residents and fellows in ACGME-accredited neurology programs were women.^9^ For woman trainees and junior faculty, attending national conferences is an opportunity to see and network with successful women in speaker and leadership roles. Such opportunities are particularly important because there is a shortage of same-sex mentorship for women in medicine.^10-12^ Specialty organizations could consider showcasing a disproportionately high representation of women in national forums to help provide role models and mitigate this shortfall in local mentorship. Perhaps the simplest way to effect this change is to give more women the authority to recruit speakers. When women are assigned leadership roles in conferences, gender equity among speakers is more likely.^13,14^

Other potential factors merit study to better explain the gender gap in rank at top-ranked academic neurology programs. Traditional measures of academic productivity such as publication quantity and impact remain important factors in promotions decisions. The lower publication rate among women is a likely reason that fewer women achieve senior academic ranks, but this is unlikely to be the sole reason.

Grant support from national funding agencies may be another factor that contributes to the gender gap in academic promotion. In a study of Canadian Institute of Health Research grants, grant application success of women when compared to men was lower when the reviewers were asked to focus on the principal investigator, rather than on the quality of the proposed science.^15^

Future studies of gender disparities in academic advancement within neurology should investigate differences in membership and leadership in national committees, as well as differences in visiting and endowed professorships. While local factors, such as administrative effectiveness, quantity and quality of teaching, mentorship, and development of curricula or educational resources may be important, these factors are harder to compare across individuals and institutions.

The strengths of our study include its large sample size and our ability to link a high proportion (83%) of speakers to the AMA Masterfile. Our study has two primary limitations. First,17% of speakers could not be matched to the AMA Masterfile, and linkage rates were lower for women than for men. While this suggests that our participation estimates may be somewhat biased, the overall high linkage rate makes it unlikely that major gender differences in speaker roles exist. Second, we limited our analysis to the five largest neurology conferences in the U.S.; it is possible that gender disparities in speaker roles are more pronounced at smaller conferences or at non-U.S. conferences. In addition, other potential drivers of disparity, such as racial and ethnic minority status, were not directly investigated.

Drivers of gender inequity in academic neurology should continue to be examined until we have probable explanations for the origins of the gender gap. Only then will our field be able to incorporate meaningful changes to allow our academic leadership to better reflect the people we train and the populations we serve. We are hopeful that with continued attention to practices that foster appropriate representation, the demographic characteristics among conference speakers will eventually reflect the population of medical school graduates entering the field of neurology, and that this will ultimately contribute to elimination of disparities in academic neurology.

## Acknowledgements

This study was funded by the Jerry Isler Neuromuscular Fund.

